# Comprehensive Transcriptome Quality Assessment Using CATS: Reference-free and Reference-based Approaches

**DOI:** 10.1101/2025.07.22.666112

**Authors:** Kristian Bodulić, Kristian Vlahoviček

## Abstract

Accurate assessment of transcriptome assembly quality is critical to ensure the reliability of subsequent transcriptomic analyses. We present CATS (Comprehensive Assessment of Transcript Sequences), a tool offering both reference-free (CATS-rf) and reference-based (CATS-rb) transcriptome quality evaluation pipelines. CATS-rf maps RNA-seq reads back to the assembled transcripts and computes four interpretable scoring components that capture common assembly errors. CATS-rb assesses transcriptome completeness via alignment to a reference genome, supporting both annotation-free and annotation-based scoring. We benchmarked CATS on 672 transcriptomes from simulated and public RNA-seq data. CATS-rf outperformed existing tools in both transcript-level accuracy assessment and demonstrated high sensitivity to diverse assembly error types. CATS-rb produced robust transcriptome completeness estimates even without external annotation, with its scoring metrics strongly reflecting assembly quality. These results highlight CATS as an accurate, interpretable, and broadly applicable framework for evaluating transcriptome assemblies.

## 1 Introduction

Accurate assembly of RNA-seq data into full-length transcripts is a prerequisite to virtually any downstream analysis procedure (1,2). Most transcriptome assemblers build de Bruijn graphs from read k-mers and reconstruct transcripts by traversing the graph paths (1–6). However, the assembly process is challenged by features intrinsic to transcriptomic data, including non-uniform coverage and alternative splicing. These issues are pronounced in low-abundance transcripts, which are frequently misassembled. De novo assemblers implement various approaches to mitigate these challenges, including read correction and normalization, multi-k-mer strategies, and clustering of isoforms into subgraphs (3–6). Benchmarking studies report considerable variability in assembly quality driven by transcriptome complexity, sequencing depth, library quality, and assembler choice. The observed variability is exacerbated by the large parameter space among assemblers, further complicating the generalization of optimal assembly strategies (7–12). These challenges underscore the need for reliable methods which assess transcriptome assembly quality and precisely characterize detected assembly errors.

Transcriptome quality assessment tools are broadly categorized by their use of reference data as reference-free or reference-based. RSEM-EVAL and TransRate are the only tools specifically performing reference-free transcriptome quality evaluation. RSEM-EVAL employs a probabilistic model that considers the joint likelihood of the assembly and the RNA-seq reads, with its performance constrained by several limitations. For instance, RSEM-EVAL’s penalty against assembly complexity may favor overly simplistic assemblies. The tool is also permissive of assemblies that merge low-coverage transcripts, leading to inflated scores for structurally implausible reconstructions. Furthermore, RSEM-EVAL’s alignment algorithm does not account for indels, reducing sensitivity to certain types of errors. The resulting transcript scores are also difficult to interpret regarding specific error types (13).

TransRate addresses some of these limitations by estimating assembly quality directly from read alignment evidence. The tool maps RNA-seq reads to assembled transcripts and uses transcript abundance to probabilistically assign multimapped reads. Transcript scores are inferred based on deviations from expected mapping patterns. Despite these strategies, TransRate often underestimates the quality of lowly expressed isoforms. Another drawback is the independent assignment of multimapped paired-reads, introducing artificial penalties for isoforms and paralogues. TransRate also poorly distinguishes alignment errors caused by low read quality or transcript variation from mapping inconsistencies resulting from misassembly (14). Additionally, the tool appears to be no longer maintained (15).

RNAQuast and SQANTI3 are widely used reference-based pipelines for transcriptome quality assessment, leveraging reference genomes to produce alignment-based and completeness metrics. Both tools rely on genomic annotations, limiting their ability to assess the fidelity of unannotated transcripts. Furthermore, completeness metrics are calculated at the isoform level, potentially introducing bias by including transcript models with overlapping exon structures. SQANTI3 is specifically designed for long-read sequencing data, making it unsuitable for evaluating short-read or hybrid transcriptome assemblies. Notably, neither tool provides a unified score reflecting overall assembly completeness, hindering direct assembly-to-assembly comparison (16,17).

Here, we introduce CATS (Comprehensive Assessment of Transcript Sequences), a framework for transcriptome quality evaluation. CATS features both reference-free (CATS-rf) and reference-based (CATS-rb) pipelines, designed to alleviate the limitations of existing methods. Both tools provide unified assembly quality scores, complemented by a comprehensive set of supporting metrics. Through extensive testing on simulated and public datasets, we demonstrate that CATS is a robust and accurate solution for transcriptome assembly assessment.

## 2 Results

### 2.1 Overview of the CATS-rf Pipeline

CATS-rf (reference-free) evaluates transcriptome assembly quality using short-read RNA-seq data employed in the assembly process (Fig. 1). The pipeline maps reads to the analysed transcripts, assigning multimapped reads probabilistically based on transcript abundance. Assembly evaluation is performed at the transcript level, integrating four score components each targeting specific assembly errors. The coverage component flags low-coverage regions suggesting unsupported insertions or redundancy, while the accuracy component detects segments exhibiting sequence inaccuracy. Both components apply penalties proportional to the length of affected regions. The local fidelity component captures inconsistent paired-read mapping within individual transcripts, suggesting structural errors such as deletions, inversions, or translocations. These inconsistencies include reads with unmapped pairs, inaccurate mapping orientations, or abnormal insert sizes. The integrity component identifies transcript fragmentation through read pairs mapping to separate transcripts, with penalties scaled based on their proximity to transcript ends. Transcript quality score is defined as the product of the four components, with the assembly quality score calculated as the mean of individual transcript scores.

**Figure 1.**
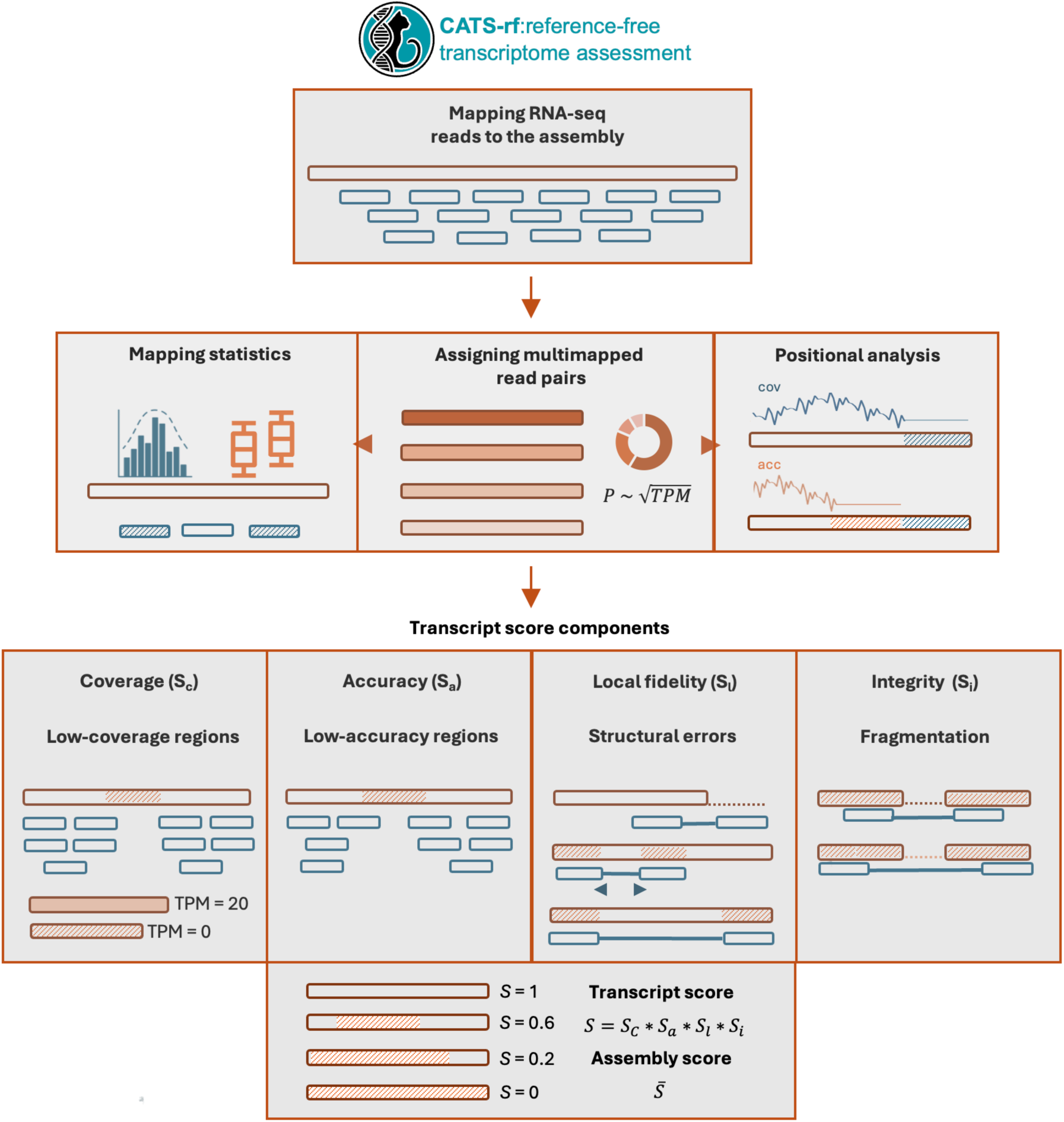
Overview of transcriptome assembly evaluation by CATS-rf. RNA-seq reads are mapped back to assembled transcripts, with multimapped reads probabilistically assigned based on transcript abundance. Read alignment data are used to compute four transcript-level score components: coverage, accuracy, local fidelity, and integrity. Each component targets a specific class of assembly errors, including insertions, mismatches, structural inconsistencies, and fragmentation. Transcript scores are calculated as the product of the four components, with the assembly score defined as the mean of individual transcript scores.

### 2.2 Association of CATS-rf Transcript Scores with Assembly Features and Quality

CATS-rf evaluation was performed on a comprehensive dataset consisting of simulated and public RNA-seq libraries (Fig. 2). We first benchmarked CATS-rf using 504 transcriptomes assembled from 126 simulated libraries, covering six model species, seven transcript coverage ranges, and three sequencing error rates. Each library was assembled with four state-of-the-art transcriptome assemblers. The distribution of CATS-rf transcript scores varied significantly across species and assemblers (Fig. 2a). Transcriptomes with higher complexity, such as those of *M. musculus and H. sapiens*, yielded lower transcript scores, with rnaSPAdes and Trinity consistently outperforming IDBA-tran and SOAPdenovo-Trans. Low coverage levels led to significantly reduced transcript scores across all assemblies (Fig. 2b). Increasing coverage beyond 5–10× generally yielded marginal improvements to transcript scores, although the gains were more evident in libraries with higher sequencing error rates.

**Figure 2.**
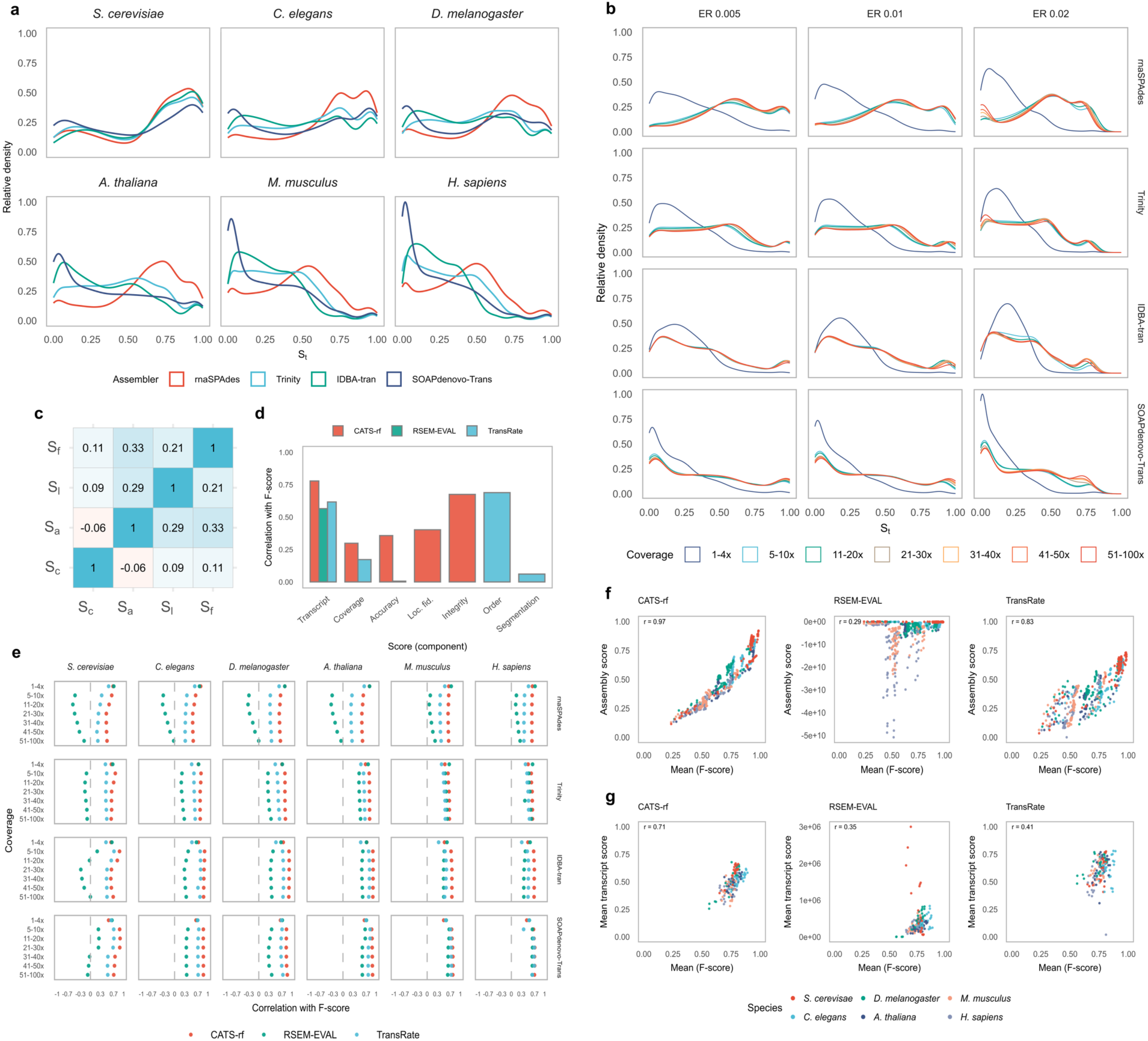
Association of CATS-rf scores with assembly features and quality. Panels a–f show results from the simulated dataset**. a** Distribution of CATS-rf transcript scores across species and transcriptome assemblers. **b** Distribution of CATS-rf transcript scores by transcript coverage, sequencing error rate, and assemblers. **c** Pairwise correlation between CATS-rf score components. **d** Correlation of CATS-rf, RSEM-EVAL, and TransRate transcript scores and score components with transcript F-scores. **e** Correlation of transcript scores from all three tools with transcript F-scores across species, coverage levels, and assemblers. **f** Correlation of assembly scores from all three tools with mean transcript F-scores. **g** Equivalent correlations (as in panel f) for the publicly available dataset. *S_t_* = transcript score, ER = error rate, *S_c_* = coverage component, *S_a_* = accuracy component, *S_l_* = local fidelity component, *S_i_* = integrity component.

CATS-rf score components were weakly inter-correlated, ranging from pairwise r=–0.06 to r=0.33 (Fig. 2c). To compare performance, transcript scores from CATS-rf, RSEM-EVAL, and TransRate were correlated with transcript F-scores derived from reciprocal best hits to reference transcripts, serving as an objective measure of accuracy (Fig. 2d). CATS-rf transcript scores exhibited the strongest correlation with F-scores (r=0.78), outperforming RSEM-EVAL (r=0.57) and TransRate (r=0.62). The CATS-rf coverage component correlated more strongly with F-scores (r=0.30) than the TransRate equivalent (r=0.17). Similarly, the CATS-rf accuracy component (r=0.36) outperformed its TransRate counterpart (r=0.01). While TransRate aggregates evidence for structural errors and fragmentation into a single “order” score, CATS-rf separates these into distinct components, both showing moderate correlations with F-scores (local fidelity: r=0.40, integrity: r=0.68).

In-depth benchmarking confirmed that CATS-rf consistently outperformed both RSEM-EVAL and TransRate across all analysed taxa and non-minimum coverage levels (Fig. 2e). The largest performance gains were found in rnaSPAdes assemblies. At the lowest coverage level, all tools exhibited similar accuracy. Additionally, we assessed the association between overall assembly scores and mean transcript F-scores (Fig. 2f). CATS-rf again outperformed its counterparts, showing stronger correlation with assembly accuracy (r=0.97, 95%CI 0.96-0.97) compared to RSEM-EVAL (r=0.29, 95%CI 0.21-0.37, z=36.14, p<0.001) and TransRate (r=0.83, 95%CI 0.80-0.86, z=17.46, p<0.001).

We also applied CATS-rf to 168 transcriptomes assembled from 42 public RNA-seq libraries. Transcript scores were generally lower than those from simulated assemblies (Extended Data Fig. 1). Consistent with simulated data, complex transcriptomes generally yielded lower scores. Among assemblers, rnaSPAdes and Trinity exhibited the highest scores. Comparative benchmark was performed on transcripts present in the corresponding reference transcriptomes (Fig. 2g). CATS-rf demonstrated the strongest correlation between mean transcript scores and mean F-scores (r=0.71, 95%CI 0.62–0.78 compared to RSEM-EVAL (r=0.35, 95%CI 0.21–0.47, z=5.14, p<0.001) and TransRate (r=0.41, 95%CI 0.28–0.53, z=4.51, p<0.001).

### 2.3 Association Between CATS-rf Transcript Scores and Assembly Errors

Next, we directly assessed the ability of CATS-rf to detect common assembly errors (Fig. 3). We introduced four increasing levels of mutations to each assembly across a subset of 12 simulated assemblies. Five mutation types were independently introduced: insertions, mismatches, deletions, fragmentation, and redundancy (cf. Methods). Mutated assemblies exhibited a significant decline in CATS-rf overall assembly score, proportional to the mutation level (Fig. 3a). An exception was observed in redundant assemblies, where the assembly score plateaued at the third redundancy level (60% of transcripts with 60-79% sequence duplication). We also analysed the coverage, accuracy, and local fidelity score components of transcripts containing increasing levels of insertions, mismatches, and deletions, respectively (Fig. 3b). All three mutation types led to significant reductions in the corresponding score components, with insertions eliciting the strongest decline.

**Figure 3.**
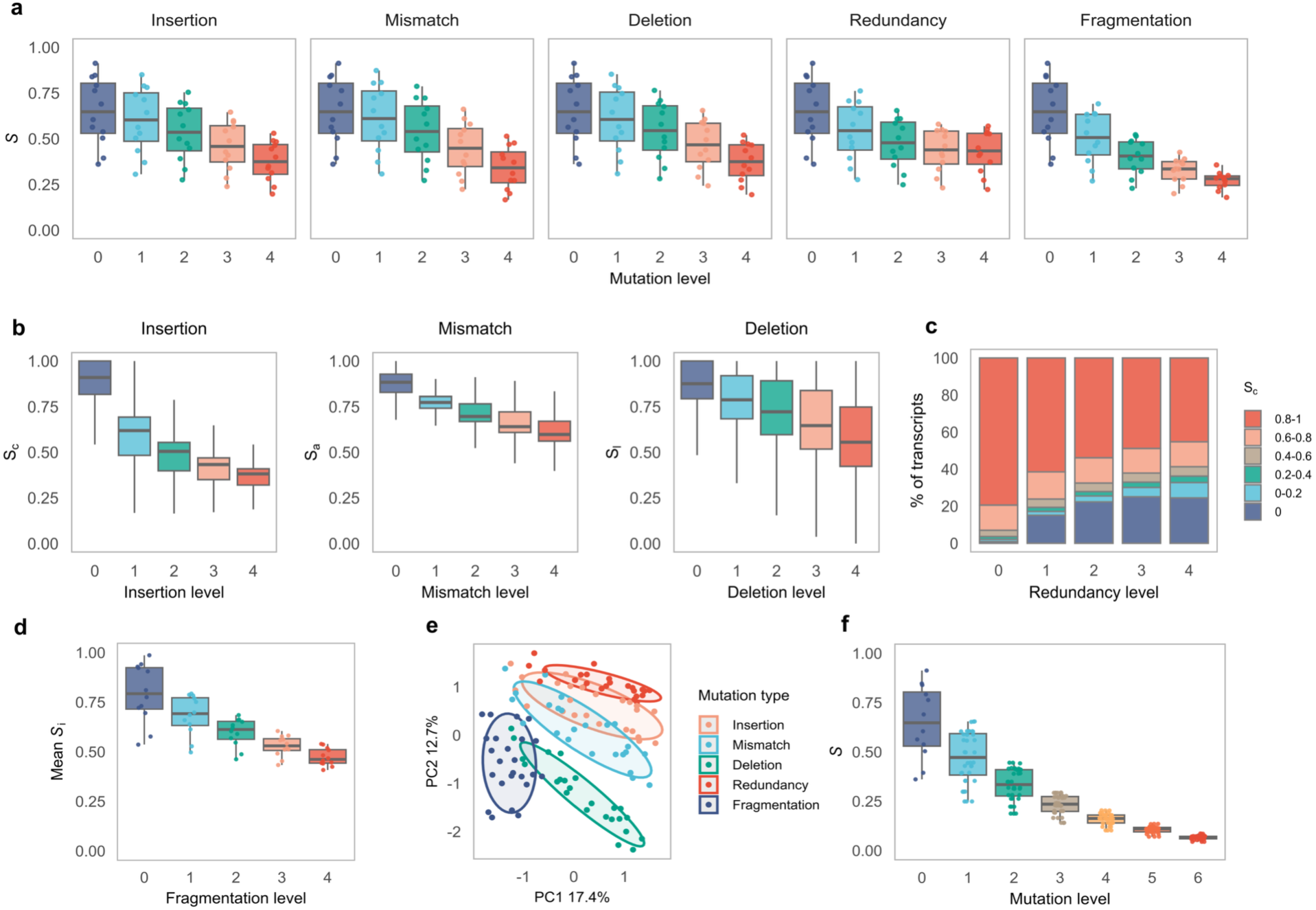
Association between CATS-rf scores and assembly errors. Panels a–e show results from transcriptome assemblies with individual mutation types. **a** Distribution of assembly scores across assemblies with increasing levels of insertions, mismatches, deletions, redundancy, and fragmentation. **b** Distribution of coverage, accuracy, and local fidelity score components in transcripts with increasing levels of insertions, mismatches, and deletions, respectively. **c** Distribution of the coverage score component across assemblies with increasing redundancy. **d** Distribution of the mean integrity score component across assemblies with increasing fragmentation. **e** Principal component analysis of mean transcript score components in assemblies with distinct mutation types. **f** Distribution of assembly scores in assemblies with increasing levels of multiplicative mutations. Boxplots represent the median and IQR of the distribution, with whiskers extending to ±1.5×IQR. Points denote individual assemblies. *S* = assembly score, *S_c_* = coverage component, *S_a_* = accuracy component, *S_l_* = local fidelity component, *S_i_* = integrity component.

Redundant assemblies exhibited a significant reduction in the coverage component. As with the overall assembly score, this decline became less pronounced after the third redundancy level (Fig. 3c). We also recorded a gradual decline of the integrity component with increasing fragmentation (Fig. 3d). To explore the scoring patterns associated with different mutation types, we performed principal component analysis (PCA) on mean score components of assemblies with all five mutation types at mutation levels 3 and 4 (Fig. 3e). Distinct clusters emerged based on mutation type, with partial overlap between assemblies containing insertions and redundancy, as well as between assemblies with insertions and mismatches.

Finally, we evaluated the performance of CATS-rf on assemblies simultaneously incorporating all mutation types (Fig. 3f). Assembly scores substantially declined with increasing mutation intensity. Moreover, the score variance decreased at higher mutation levels.

### 2.4 Overview of the CATS-rb Pipeline

CATS-rb (reference-based) evaluates transcriptome assemblies by aligning transcripts to the reference genome of the corresponding species (Fig. 4). The pipeline calculates a broad range of metrics, including transcript mappability, exon statistics, isoform composition, and structural consistency. Transcripts are classified as structurally inconsistent if they exhibit a low alignment proportion or if their segments map to disjunct genomic regions.

**Figure 4.**
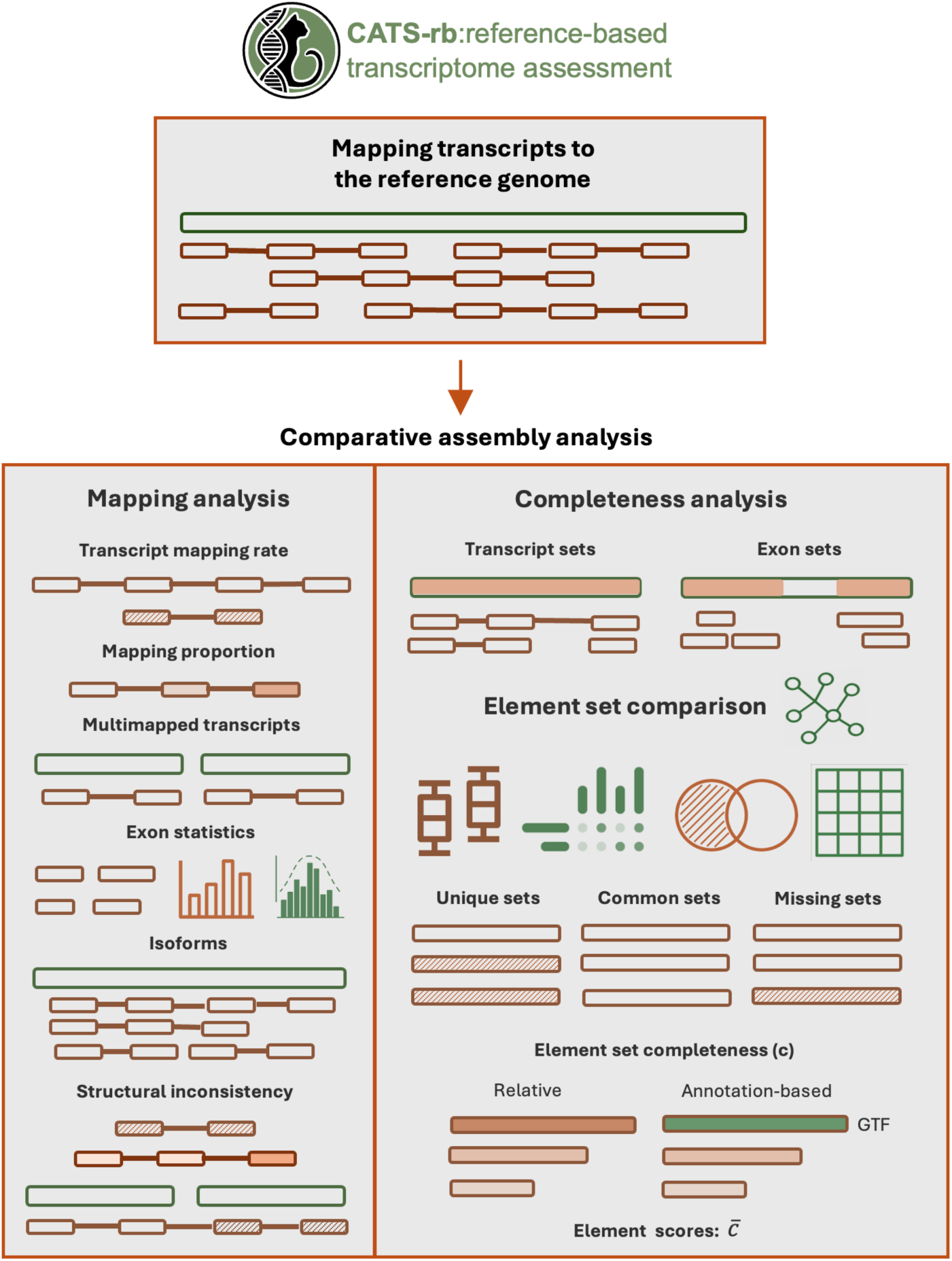
Overview of transcriptome assembly evaluation by CATS-rb. Transcripts are mapped to the reference genome and several mapping metrics are evaluated, including mappability, exon-level statistics, and structural consistency. Completeness analysis is performed across two categories of element sets, determined by overlapping transcript or exon genomic coordinates within each transcriptome assembly. Element set coordinates are overlapped between the analysed transcriptomes, constructing a graph where edges represent element set overlaps and connected components denote element set groups. In-depth analysis of element sets is performed, visualized using UpSet plots, Venn diagrams, and hierarchical clustering heatmaps. Element set completeness is defined as the relative length of each set compared to the longest set within its group. The analysis can be extended using reference element sets derived from genomic annotation, with assembly set groups defined by shared overlaps with reference sets. Transcript and exon scores are calculated as the mean completeness of the respective element sets.

The core functionality of CATS-rb lies in the assessment of assembly completeness. Briefly, completeness analysis is based on two types of non-redundant element sets, derived by merging overlapping genomic coordinates of transcripts or exons within each transcriptome assembly. Undirected graphs are independently constructed for transcript and exon sets, with edges indicating coordinate overlaps between sets across transcriptomes. In each graph, overlapping element sets are grouped into connected components, with the longest set designated as the representative. The completeness of each set is quantified as its length relative to the representative set in the corresponding group. Relative transcript and exon scores are computed as the mean completeness of the respective element sets, providing complementary insights into assembly quality. The transcript score captures large-scale structural issues, such as missing isoforms and fragmentation, while the exon score detects partial incompleteness and localized errors.

CATS-rb is primarily designed to estimate assembly completeness relative to the supplied transcriptome assemblies. However, the analysis can be supplemented using non-redundant reference element sets derived from genomic annotation. In this mode, assembly element set groups are defined according to shared overlaps with reference sets. Annotation-based element scores are calculated analogously to the relative scores, providing absolute completeness measures.

### 2.5 CATS-rb Performance Assessment

CATS-rb benchmark was performed on an extensive dataset of simulated and public RNA-seq libraries (Fig. 5), previously used in CATS-rf evaluation. We first examined the general features of CATS-rb metrics using the simulated dataset. Relative transcript and exon scores varied substantially across assemblers and species, with complex transcriptomes generally showing worse performance (Fig. 5a, Extended Data Fig. 2). Consistent with the reference-free pipeline, rnaSPAdes and Trinity outperformed IDBA-tran and SOAPdenovo-Trans. Reference assemblies yielded consistently high relative scores (transcript: 0.95-0.99, exon: 0.97-0.99). Assemblies generated at the lowest coverage levels showed the worst performance, with transcript and exon scores typically plateauing beyond 5–10× coverage (Fig. 5b, Extended Data Fig. 3). However, assemblies with the highest sequencing error rate continued to significantly improve with increasing depth.

**Figure 5.**
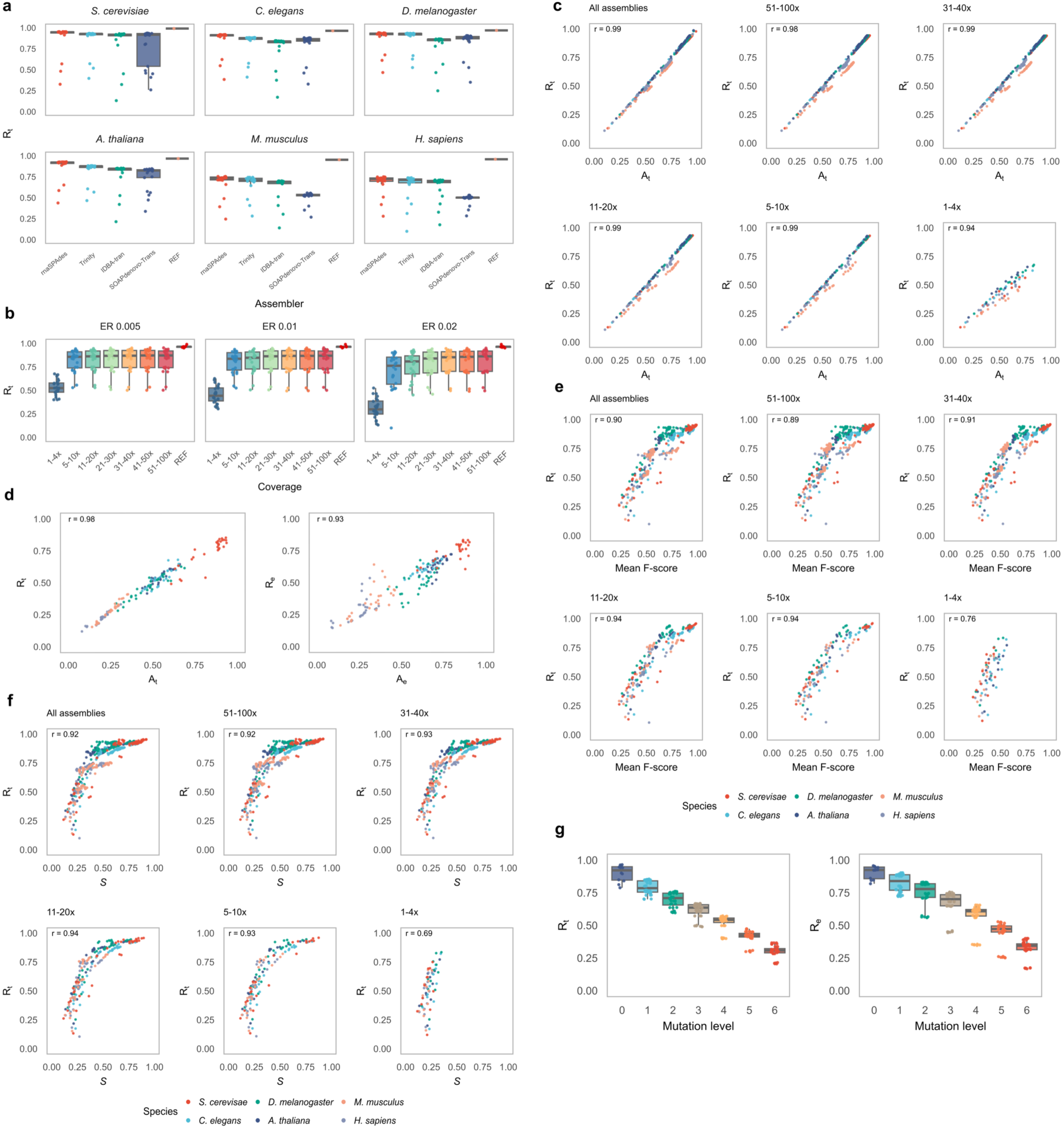
Performance assessment of CATS-rb. **a** Distribution of relative transcript scores across species and assemblers. **b** Distribution of relative transcript scores by transcript coverage and sequencing error rate. **c** Correlation between relative and annotation-based transcript scores in the full dataset and five subsets of simulated assemblies with decreasing coverage, as indicated in subplot titles. **d** Equivalent correlations (as in panel c) in the publicly available dataset. **e** Correlation between relative transcript scores and mean transcript F-scores across the full dataset and assembly subsets with decreasing coverage. **f** Correlation between relative transcript scores and CATS-rf assembly scores across the full dataset and assembly subsets with decreasing coverage. **g** Distribution of relative scores in assemblies with increasing levels of multiplicative mutations. Boxplots represent the median and IQR of the distribution, with whiskers extending to ±1.5×IQR. *R_t_* = relative transcript score, REF = reference transcriptome, ER = error rate, *A_t_* = annotation-based transcript score, *R_e_* = relative exon score*, A_e_* = relative annotation-based exon score*, S* = CATS-rf assembly score.

CATS-rb relative scores were evaluated across the complete dataset and five assembly subsets with decreasing coverage for each species. The complete dataset included the reference transcriptome and all simulated assemblies. In contrast, each subset comprised only simulated assemblies and was defined by the coverage range of its highest-coverage assemblies, ranging from 51–100× to 1–4×. We observed a strong correlation between relative and annotation-based scores across the full dataset and all subsets (Fig. 5c, Extended Data Fig. 4). The stated association was strong even in the lowest-coverage subset (transcript: r=0.94, 95%CI 0.90-0.96, exon: r=0.95, 95%CI 0.93-0.97). These findings were further validated using assemblies from public libraries, where separate CATS-rb runs were conducted on datasets comprising the reference transcriptome and assemblies from each library (Fig. 5d). The public dataset displayed similarly strong correlations for transcript (r=0.98, 95%CI 0.97–0.98) and exon (r=0.93, 95%CI 0.91–0.95) scores.

The accuracy of relative transcript scores was further demonstrated by their strong association with mean transcript F-scores across the full dataset (r=0.90, 95%CI 0.88–0.91) and all subsets (Fig. 5e). Analogous results were observed for relative exon scores (Extended Data Fig. 5). Even in the lowest-coverage subset, observed correlations remained strong for both transcript (r=0.76, 95%CI 0.64–0.85) and exon (r=0.79, 95%CI 0.68–0.86) scores. We also analysed the relationship between CATS-rb and CATS-rf quality estimates (Fig. 5f). A strong correlation between CATS-rb relative transcript scores and CATS-rf assembly scores was observed in the full dataset (r=0.92, 95%CI 0.91-0.93) and in all assembly subsets, including the lowest-coverage subset (r=0.69, 95%CI 0.54-0.79). Similar trends were observed for exon scores (Extended Data Fig. 6). In the lowest-coverage subset, CATS-rb scores were notably higher than both the mean transcript F-scores and the CATS-rf assembly scores.

To further evaluate its performance, CATS-rb was applied to assembly groups including native and mutated assemblies, the latter comprising six levels of multiplicative mutations. CATS-rb successfully captured the introduced errors, as reflected by a gradual decline in relative scores with increasing mutation levels (Fig. 5g).

Lastly, we assessed the performance of CATS-rb in detecting chimerism by artificially merging transcripts from reference assemblies (Extended Data Fig. 7). CATS-rb demonstrated strong performance in classifying chimerism as a structural inconsistency, exhibiting high sensitivity (93.5%-96.8%) and specificity (97.0%-99.8%) across all assemblies.

## 3 Discussion

This study presents CATS, a framework for transcriptome assembly evaluation combining reference-free (CATS-rf) and reference-based (CATS-rb) approaches. CATS-rf leverages short-read RNA-seq mappings to detect specific transcript errors through interpretable score components. While validated using Illumina data, CATS-rf is compatible with all short-read sequencing platforms. In contrast, CATS-rb evaluates assemblies by aligning transcripts to a reference genome, providing robust completeness scores without relying on genomic annotation. CATS-rb does not require input RNA-seq reads, making it applicable to any sequencing technology.

CATS-rf demonstrated superior performance compared to existing reference-free transcriptome assessment tools across a broad range of simulated libraries. While CATS-rf continued to significantly outperform RSEM-EVAL and TransRate on public datasets, quality estimates of all tools exhibited weaker correlations with transcript accuracy. This reduction may result from the alignment-based filtering strategy applied to public datasets, which aimed to retain only transcripts represented in the reference transcriptome. Such filtering may have excluded poorly assembled transcripts that failed to meet alignment thresholds, while retaining biologically relevant features absent from the reference transcriptome, such as unannotated splice variants and transcript fusions. These effects likely biased the estimation of true transcript accuracy on public assemblies. Furthermore, the reduced correlation may also reflect greater variability in coverage between public and simulated datasets.

The scoring scheme of CATS-rf provides high sensitivity to distinct assembly errors, as evidenced by a systematic reduction of CATS-rf scores in assemblies with increasing levels of several mutation types. Moreover, CATS-rf demonstrated better performance in detecting assembly errors compared to TransRate, as reflected by stronger correlations between its score components and transcript accuracy. This improvement stems from the design of CATS-rf score components, which effectively separate mapping inconsistencies due to sequencing noise and transcript variation from those caused by misassembly. Notably, evidence associated with each component cannot be specifically assigned to a single misassembly event, as multiple error types can contribute to the observed inconsistencies. However, low correlations between CATS-rf score components, along with distinct clustering of error types in PCA, suggest that different error classes yield characteristic score component distributions.

While CATS-rf score components penalize unsupported fusion sites, CATS-rf does not directly detect chimerism. This limitation is likely minor, as studies report a low incidence of chimeric artifacts produced by modern assemblers (10,12,18). In contrast, TransRate computes a chimerism score based on the uniform coverage assumption (14), which is unrealistic due to fragmentation and GC content biases (19–21). Meanwhile, the reference-based approach employed in CATS-rb demonstrated strong performance in identifying chimerism as a form of structural inconsistency. However, the obtained metrics should be interpreted in the context of rare-event classification.

CATS-rb is the first tool that directly estimates transcriptome assembly completeness without relying on reference transcriptomes or genomic annotation, making it suitable for the analysis of poorly annotated organisms. Evaluation on simulated and public datasets revealed an exceptionally strong correlation between CATS-rb relative and annotation-based scores. Moreover, CATS-rb scores were closely associated with F-scores and CATS-rf assembly scores, further validating the accuracy of both tools. While this association remained strong across all coverage levels, the absolute values of CATS-rb scores significantly differed from F-scores and CATS-rf scores at the lowest coverage. This discrepancy likely reflects the differing evaluation scope: both CATS-rf and F-scores assess transcript-level accuracy, whereas CATS-rb captures assembly completeness. As such, joint interpretation of CATS-rf and CATS-rb scores offers a complete representation of assembly quality.

CATS-rb showed high sensitivity to assembly errors, as reflected by significantly reduced scores in mutated assemblies. Additionally, reference transcriptomes consistently received near-maximum scores, suggesting minimal score inflation. These findings indicate that the tool effectively estimates transcript completeness on complex and low-quality datasets. As such, we propose that CATS-rb is valuable in three scenarios: evaluating assemblies generated from the same library, comparing transcriptomic content across libraries, and assessing absolute completeness using a reference transcriptome or genomic annotation.

The performance of both tools within the CATS framework is intrinsically tied to their respective alignment strategies. CATS-rf utilizes Bowtie2 with optimized parameters that balance between error detection and mapping stringency. In parallel, CATS-rb leverages Spaln for robust transcript-to-genome alignment, incorporating species-specific parameters that can be user-adjusted. While we do not claim these aligners are universally optimal, both are regarded as highly accurate (22,23), and the selected parameters have been specifically tuned for transcriptome assembly evaluation.

Benchmarking both tools revealed substantial variation in assembly quality across species, with complex transcriptomes consistently yielding lower scores. This trend reflects the assembly challenges posed by isoform diversity, paralogous gene families, and repetitive sequences (1,7,9). Both tools ranked rnaSPAdes and Trinity above IDBA-tran and SOAPdenovo-Trans, in agreement with recent benchmarking studies (7,9,11,12). However, this result should be validated using assemblies generated with a broader range of k-mer lengths. Increasing sequencing depth beyond a certain threshold did not significantly improve assembly scores, particularly in low-error datasets. These results are consistent with studies highlighting the diminishing returns of deeper sequencing for accurate transcript reconstruction (19,24).

In summary, CATS provides a comprehensive framework for evaluating transcriptome assemblies, combining the strengths of reference-free and reference-based approaches. Its consistent performance across diverse conditions makes it well-suited for a broad range of transcriptomic studies. By offering interpretable and robust metrics, CATS facilitates reliable quality control of transcriptome assemblies.

## 4 Methods

### 4.1 CATS-rf

#### 4.1.1 Read Mapping

CATS-rf generates transcript-level quality metrics using short-read RNA-seq data employed in the transcriptome assembly process. The pipeline begins by mapping the reads to the evaluated assembly using Bowtie2 (22), with parameters optimized for accurate detection of assembly errors. Read alignment is performed in global mode using the sensitive preset. To accommodate transcripts with multiple isoforms, the default maximum number of alignments per read is set to 10. A mismatch penalty of 2 is applied to allow the detection of potentially inaccurate transcript regions, while controlling for false-positive alignments. Secondary alignments with edit distance exceeding a defined threshold are discarded, reducing off-target mappings. This threshold is set to the sum of the maximum edit distance of the read and 10% of mean read length.

Reads that map to multiple transcripts are probabilistically assigned to a single transcript based on expression levels calculated by Kallisto using default parameters (25). Specifically, the probability *P* of assigning a read to transcript *t* is proportional to transcript abundance, measured in transcripts per million (TPM)

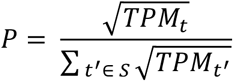

where *S* is the set of all candidate transcripts *t’* for a given read.

The introduced transformation of TPM aims to increase the effective number of assigned reads in lowly-expressed transcripts. To ensure mapping consistency, CATS-rf enforces co-assignment of paired reads. If read pairs map to different transcripts, both are reassigned to the most abundant shared transcript. This eliminates artifacts that could bias downstream scoring components based on pair mapping.

#### 4.1.2 Transcript Score Components

CATS-rf computes transcript-level quality scores by analyzing per-base coverage patterns and the consistency of paired-read mappings. Per-base transcript statistics are extracted from read mappings using pysamstats (26). Four score components are defined, each corresponding to distinct types of assembly errors.

The coverage score component *S_c_* penalizes transcript regions with low read coverage, primarily aiming to capture unsupported insertions and redundancy. Low coverage regions (LCR) are identified as rolling windows of size *L_c_*, where the mean read depth falls below a defined threshold. Each LCR is assigned a coverage penalty *P_c_* calculated as:

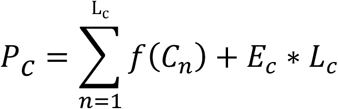

Here, *C_n_* denotes the coverage of the n_th_ base. *E_c_* represents the LCR extension penalty, and *f(C_n_)* is a function of per-base coverage defined as:

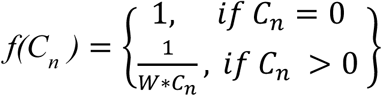

where *W* is the base coverage weight.

The coverage component is defined as the complement of the normalized sum of coverage penalties:

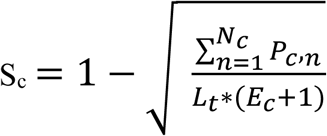

where *N_c_* is the number of LCRs per transcript and *L_t_* denotes transcript length.

By definition, uncovered transcripts are assigned a coverage component of zero with other score components remaining undefined.

The accuracy score component *S_a_* identifies regions exhibiting inaccurate transcript sequence. Accuracy is calculated as the fraction of aligned read bases matching the transcript base. Low-accuracy regions (LAR) are defined among covered bases as rolling windows of size *L_a_* with mean accuracy below a specific threshold. The definition is refined by the exclusion of fully accurate bases on LAR ends. Each LAR is assigned an accuracy penalty *P_a_* given by:

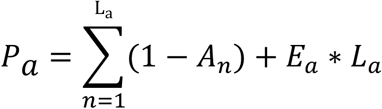

where *A_n_* represents the accuracy of the n_th_ base and *E_a_* denotes the LAR extension penalty.

The accuracy component is defined as the complement of the normalized sum of accuracy penalties:

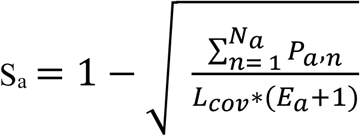

where *N_a_* is the number of LARs per transcript and *L_cov_* denotes the number of covered transcript bases.

If a paired-end library is provided, CATS-rf calculates the local fidelity and integrity score components based on paired-read mapping patterns. The local fidelity score component *S_l_* detects mapping inconsistencies of read pairs within individual transcripts, capturing structural errors such as deletions, inversions, or translocations. This component incorporates three transcript-level metrics: number of reads whose pair fails to map to the assembly (*N_u_*), number of pairs mapping to the same transcript in unexpected orientations (*N_o_*), and total transcript distance penalty. Unexpected orientations are identified based on library strandness, which can be supplied by the user or automatically detected.

The distance penalty *P_d_* is computed for each properly oriented pair mapped to the same transcript using a segmented function of the observed pair distance *d*:

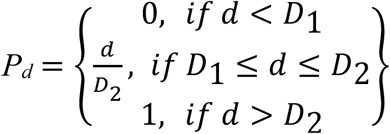

where *D_1_* and *D_2_* denote distance outlier thresholds defined as:

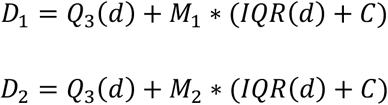

Here, *Q_3_(d)* is the third quartile and *IQR(d)* is the interquartile range of the observed pair distance. *M_1_* and *M_2_* multiplicative factors reduce the impact of false-positive outliers, which may result from variations in library preparation. The correction factor *C* ensures threshold robustness in libraries with a high proportion of overlapping read pairs.

These metrics are combined into the local fidelity component:

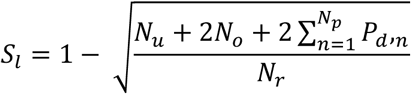

where *N_p_* refers to the number of properly oriented pairs mapped to the same transcript and *N_r_* is the total number of mapped reads per transcript.

The integrity score component *S_i_* captures transcript fragmentation by detecting bridging events, defined as read pairs mapped to separate transcripts. For each bridging event, a bridge penalty *P_b_* is computed as the mean of the relative distances of each read from the center of its respective transcript. The distances are normalized such that a value of 0 corresponds to the transcript center and a value of 1 to the transcript ends. Bridge index *B* for a transcript is then calculated as:

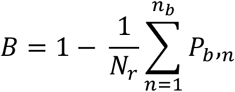

where *n_b_* is the number of bridging events per transcript.

The integrity component is defined by a sigmoid transformation of the bridge index:

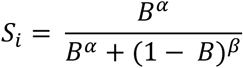

Here, *α* and *β* denote tunable compression factors that regulate sensitivity to fragmentation patterns. The transformation compensates for consistent mapping of read pairs to accurately assembled interior transcript regions, inherently weakening the fragmentation signal.

The transcript score *S_t_* is calculated as the product of the defined score components:

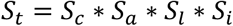

The assembly score *S* is calculated as the mean of individual transcript scores:

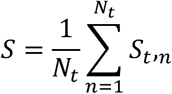

where *N_t_* is the total number of transcripts. The assembly score provides a normalized estimate of assembly quality, equally weighting distinct assembly error types.

In addition to transcript scores, CATS-rf computes and visualizes a comprehensive set of mapping and positional metrics to support quality assessment. These include coverage and accuracy statistics such as the number of uncovered and inaccurate bases, mean transcript coverage and accuracy, per-base and positional distributions of coverage and accuracy, as well as length distributions of LCRs and LARs. Paired-read mapping metrics encompass the number of inconsistently mapped read pairs, bridging events, and potentially fragmented transcripts.

### 4.2 CATS-rb

#### 4.2.1 Transcriptome Assembly Mapping

CATS-rb performs reference-based transcriptome assembly assessment by aligning transcripts to a reference genome using Spaln, a highly accurate spliced aligner (23). The pipeline comprises three core tools that perform genome index generation, transcriptome mapping, and comparative mapping analysis. During index generation, users can define the maximum gene size. Mapping parameters are highly customizable to accommodate assembly-specific features, including library strandness, species-specific presets, maximum number of alignments per transcript, minimum intron size, splice site recognition criteria, and weighting of alignment scores based on coding potential and translation signals.

#### 4.2.2 Comparative Transcriptome Assembly Analysis

CATS-rb workflow evaluates assemblies in terms of accuracy and completeness. The analysis begins by processing exon-level alignment output from Spaln, which is used to reconstruct transcript-level mappings. Exons originating from the same transcript are assigned to separate transcript mappings under the following conditions: 1) the exon overlaps the preceding exon in transcript coordinates by more than 50% of the preceding exon’s length; 2) the exon maps to a different genomic scaffold or strand than the preceding exon; or 3) the exon maps to the same scaffold and strand, but the distance to the preceding exon exceeds the intron length threshold.

The pipeline evaluates multiple aspects of assembly mapping, including transcript mappability, multimapping, exon count per transcript, exon length distribution, isoform representation, and structural consistency. A transcript is classified as structurally inconsistent if its alignment proportion falls below a predefined threshold or if it contains regions that map to disjunct genomic loci. The latter is assessed by identifying transcript segments that overlap by less than a specified proportion of their lengths and map either to different scaffolds, to opposite strands on the same scaffold, or to distant regions on the same scaffold and strand (beyond the intron length threshold). Flagged pairs are excluded if one segment is shorter than 10% of the other segment’s length. This definition is further refined by removing cases where a separately mapped transcript segment covers more than 90% of each segment in the marked pair (in transcript coordinates). The described structural consistency criteria are primarily designed to detect chimerism, while also capturing events resulting in significant mismatches to the reference genome, arising either from assembly errors or genuine biological variation.

The primary contribution of CATS-rb is the analysis of assembly completeness, which introduces two types of element sets as units for transcriptome assembly comparison. Precisely, transcript and exon sets are defined by merging the overlapping transcript or exon genomic coordinates within a given transcriptome into non-redundant segments. For multimapped transcripts, element sets are constructed using the most accurate mapping, with ties resolved by selecting the matching mapping across the analysed transcriptomes. CATS-rb builds separate undirected graphs for both element set types, in which vertices represent element sets and edges indicate overlaps between sets of the compared assemblies. Overlaps are determined using a predefined overlap length threshold for edge specification. Element set groups are defined as connected graph components. If multiple sets from the same assembly fall within a single group, only the longest is retained. The group representative is defined as the longest set in the group. Element set completeness *c* is calculated as the length of the set relative to the representative set of its group. Sets absent from a group are assigned a completeness of 0. The relative element score *R* for each assembly is computed as the mean element set completeness across all groups:

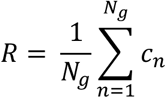

Here, *N_g_* denotes the number of element set groups. This procedure is applied independently to transcript and exon sets to produce relative transcript and exon scores.

The completeness assessment also provides a detailed evaluation of element set distribution across the compared transcriptome assemblies. This includes an in-depth analysis of missing, common, and unique element sets, along with element set UpSet plots and Venn diagrams. Assemblies are also hierarchically clustered based on element set completeness using complete linkage clustering with the Euclidean distance metric.

The presented method evaluates assembly completeness relative to the compared transcriptomes without requiring external genomic annotation. Additionally, CATS-rb can perform an annotation-based analysis using non-redundant reference element sets. These sets are constructed by separately merging overlapping transcript coordinates and overlapping exon coordinates defined by external genomic annotation. The annotation-based workflow follows the same principles as relative completeness evaluation, while grouping transcriptome assembly element sets based on shared overlaps with reference sets. As such, reference sets are considered the representative for each assembly set group. Annotation-based element scores *A* are calculated analogously to relative element scores, providing an absolute metric of assembly completeness.

Genomic annotation is also used to calculate element set sensitivity and specificity for each transcriptome assembly. Sensitivity is defined as the proportion of reference sets matched by assembly element sets, while specificity reflects the proportion of assembly element sets matched by reference sets. A set is considered matched if its genomic coordinates cover more than a predefined fraction of the corresponding set’s length.

### 4.3 Performance Evaluation of CATS

#### 4.3.1 RNA-seq Library Preparation, Transcriptome Assembly, and Mutation Simulation

We first performed an extensive evaluation of CATS performance using a diverse set of RNA-seq libraries simulated with the Polyester R package (version 1.32.0) (27). A total of 126 paired-end, unstranded libraries were simulated from coding and non-coding transcripts of six species*: Saccharomyces cerevisiae, Caenorhabditis elegans*, *Drosophila melanogaster*, *Arabidopsis thaliana*, *Mus musculus,* and *Homo sapiens* (Supplementary Table 1). For each species, 21 libraries were generated by combining seven transcript coverage levels with three sequencing error rates. Coverage levels were simulated by randomly sampling transcript coverages from the following ranges: 1–4x, 5–10x, 11–20x, 21–30x, 31–40x, 41–50x, and 51– 100x. A uniform error distribution model was applied, with error rates set to 0.005, 0.01, or 0.02 for each coverage level. Reads were 100 bp in length, and fragments were generated using an RNA fragmentation protocol with empirical fragment length distribution provided by Polyester. In addition to the simulated dataset, we benchmarked CATS on 42 publicly available RNA-seq libraries from the same six species, obtained from the Sequence Read Archive (SRA) database (Supplementary Table 2). Adapter sequences were removed and low-quality bases trimmed from all public libraries using Trim Galore (version 0.6.1) with the parameters --stringency 5 -q 5 (28). Transcriptomes from both simulated and public libraries were assembled using four de novo assemblers: rnaSPAdes (version 3.15.4), Trinity (version 2.15.1), IDBA-tran (version 1.1.1), and SOAPdenovo-Trans (version 1.0.4), each with default parameters. This resulted in a total of 504 simulated assemblies and 168 assemblies generated from public libraries.

We also tested the ability of CATS to detect assembly errors. Firstly, we independently introduced varying levels of mutations into a subset of simulated assemblies, including insertions, mismatches, deletions, redundancy, and fragmentation. Mutations were simulated in 12 assemblies generated using rnaSPAdes and Trinity, derived from libraries representing the six earlier stated species, each with transcript coverage range of 21–30× and a sequencing error rate of 0.005. For every mutation type, four levels of severity were simulated. In the case of insertions, mismatches, and deletions, a single mutated region was introduced at a random position within the selected transcripts. The proportion of affected transcripts was set to 15%, 30%, 45%, and 60%, with the length of the mutated regions scaled accordingly (15%, 30%, 45%, or 60% of the transcript length). For mismatches, each base within the mutated region was assigned a 25% probability of substitution, with equal likelihood for each alternative base. Redundancy was simulated by duplicating a corresponding transcript subsequence beginning from the 5’ transcript end, with 20%, 40%, 60%, and 80% of transcripts containing duplicated regions covering 20-39%, 40-59%, 60-79%, and 80-99% of the transcript length, respectively. Fragmentation was simulated by splitting transcripts at a random position between the 40th and 60th percentiles of transcript length, affecting 20%, 40%, 60%, and 80% of transcripts, respectively.

In addition to isolated mutation types, we performed a multiplicative error simulation, applying all five mutation types sequentially to the same 12 assemblies. The mutations were applied in the following order: deletion, mismatch, insertion, fragmentation, and redundancy. The simulation was conducted at six mutation severity levels (10%, 20%, 30%, 40%, 50%, and 60%), which controlled both the proportion of affected transcripts and the length of the affected region. To introduce stochastic variation, the multiplicative simulation was applied three times to each assembly. All mutation simulations were performed using custom R scripts.

#### 4.3.2 Performance Evaluation of CATS-rf

To compare the performances of CATS-rf, RSEM-EVAL (version 1.11) and TransRate (version 1.0.4), we ran each tool on all simulated and public assemblies using their default parameters. For simulated assemblies, the fragment length distribution mean required by RSEM-EVAL was set to 180 bp, in accordance with the empirical distribution provided by Polyester. For public assemblies, fragment length distribution parameters were estimated using Kallisto (version 0.50.1). TransRate was obtained from a Github repository that addressed inaccurate results produced by the current official release (29). To evaluate transcript-level accuracy, we used CRB-BLAST (30) to identify reciprocal best alignments between each tested and reference transcriptome. For each aligned transcript pair, we calculated the F-score as the harmonic mean of precision and recall. Precision was calculated as the number of aligned bases relative to the tested transcript length, while recall was defined as the number of aligned bases relative to the reference transcript length. Tested transcripts without hits were assigned an F-score of 0. For assemblies generated from public libraries, CRB-BLAST was applied exclusively to transcripts represented in the reference transcriptomes. These transcripts were identified using BLAT with sensitive parameters (-tileSize=8 -maxIntron=0) (31), retaining only alignments with F-score greater than 0.5.

The performance of CATS-rf, RSEM-EVAL, and TransRate was evaluated with the Spearman correlation coefficient between transcript quality scores and F-scores, as well as between the assembly scores and the mean F-score. Assembly-level correlations were compared using Meng’s z-test for overlapping correlations (R package cocor, version 1.1.4) (32), with confidence intervals for Spearman’s rho calculated using Fisher’s z-transformation (R package DescTools, version 0.99.60) (33). Additionally, we assessed the ability of CATS-rf to capture common assembly errors by comparing CATS-rf assembly scores and transcript-level score components across assemblies with increasing levels of simulated mutations. CATS-rf performance assessment was performed using custom R scripts.

#### 4.3.3 Performance Evaluation of CATS-rb

The performance of CATS-rb was also evaluated using both simulated transcriptome assemblies and assemblies generated from public RNA-seq libraries. Reference genomes and annotations were obtained from Ensembl (Supplementary Table 3). Genome indexing, transcriptome mapping, and mapping comparison were performed using default CATS-rb parameters, with adjustments to maximum gene sizes and species-specific parameters (Supplementary Table 4). CATS-rb was first applied to datasets comprising each species’ reference transcriptome and the complete set of simulated assemblies. Relative transcript and exon scores were analysed for their association with library characteristics.

We further examined the robustness of relative transcript and exon scores by running CATS-rb on five assembly subsets. The subsets excluded reference transcriptomes and consisted of simulated assemblies with progressively lower transcript coverage. Each subset was defined by the coverage range of its highest-coverage assemblies: 51–100x, 31–40x, 11–20x, 5–10x, and 1–4x. Robustness of relative scores was assessed by examining their correlation with annotation-based scores, mean transcript F-scores, and CATS-rf assembly scores. Relative and annotation-based score correlations were also evaluated on the public datasets, with separate CATS-rb runs for each library alongside the corresponding reference transcriptome.

We also assessed the sensitivity of CATS-rb to assembly errors by testing it on assemblies subjected to multiplicative mutation simulation. Each tested set included the native assembly and triplicates of assemblies generated with six increasing levels of mutation intensity.

To evaluate the ability of CATS-rb to classify chimerism as a structural inconsistency, we performed chimerism simulation by randomly merging transcript pairs from the reference transcriptomes of the six analysed taxa into single transcripts. Prior to merging, random deletions were introduced at fusion sites of both transcripts, affecting up to 3% of each transcript length. The resulting assemblies contained 10% chimeric transcripts. CATS-rb performance evaluation was conducted using custom R scripts.

## 5 Code Availability

The CATS framework is implemented in R and Bash. The source code is available in the following GitHub repositories:

CATS-rf: https://github.com/bodulic/CATS-rf

CATS-rb: https://github.com/bodulic/CATS-rb

Both repositories are distributed under the MIT license.

In addition, the code used to perform the CATS benchmark is available at: https://github.com/bodulic/CATS_benchmark

## 6 Data Availability

CATS benchmark results used for figure generation are provided as stand-alone tables and are publicly accessible via the following GitHub repository: https://github.com/bodulic/CATS_benchmark

## Supporting information

Supplementary Data Description

## 7 Acknowledgments

We would like to thank Karlo Jambrošić for designing the CATS logo.

## 9 Author contributions

Conceptualization, K.B., K.V.; data curation, K.B.; formal analysis, K.B.; investigation, K.B.; methodology, K.B.; project administration, K.V.; resources, K.V.; software, K.B.; supervision, K.V.; validation, K.B., K.V.; visualization, K.B.; writing of the original draft, K.B.; writing review and editing, K.V.

