## Supplementary figures and images for "Comprehensive Transcriptome Quality Assessment Using CATS: Reference-free and Reference-based Approaches"

### Extended Data Figures

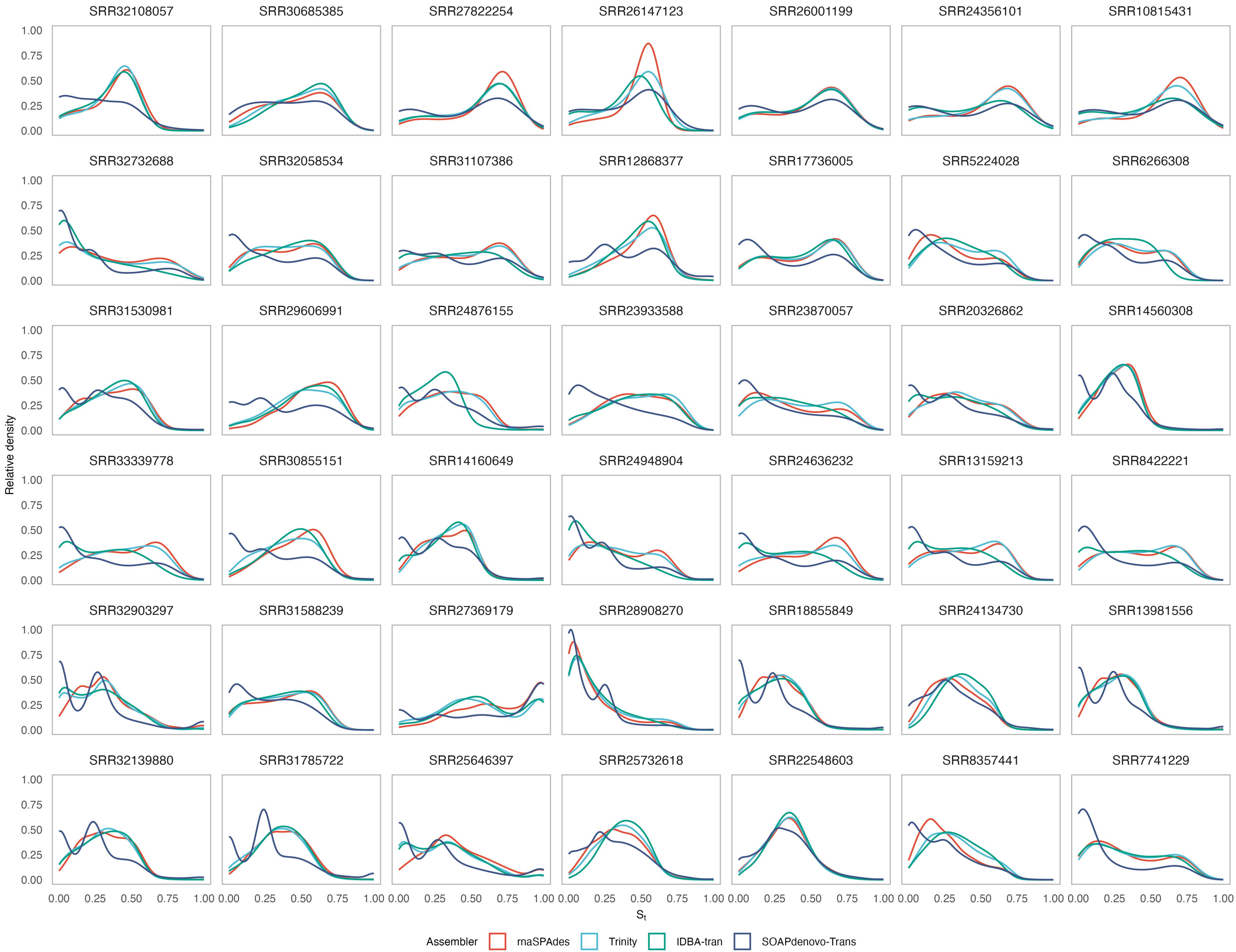

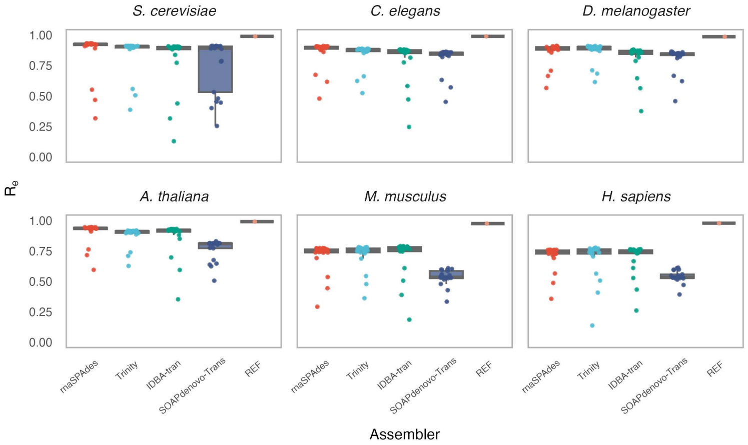

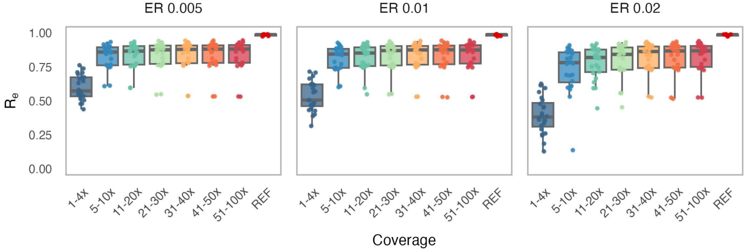

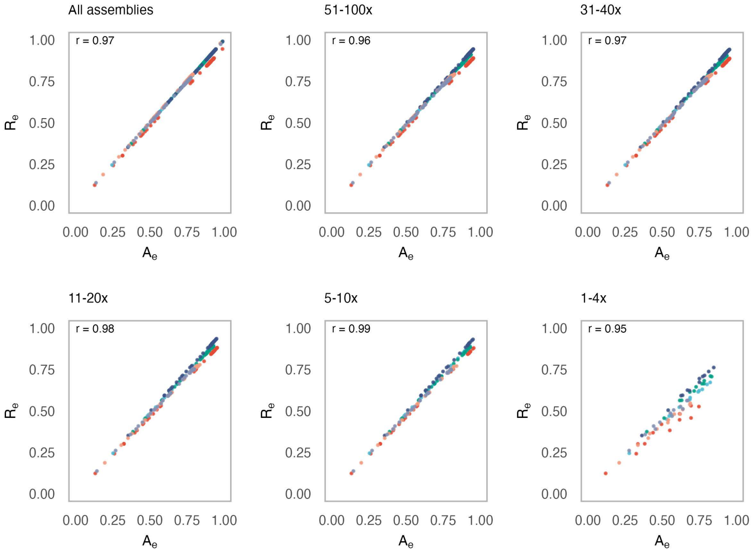

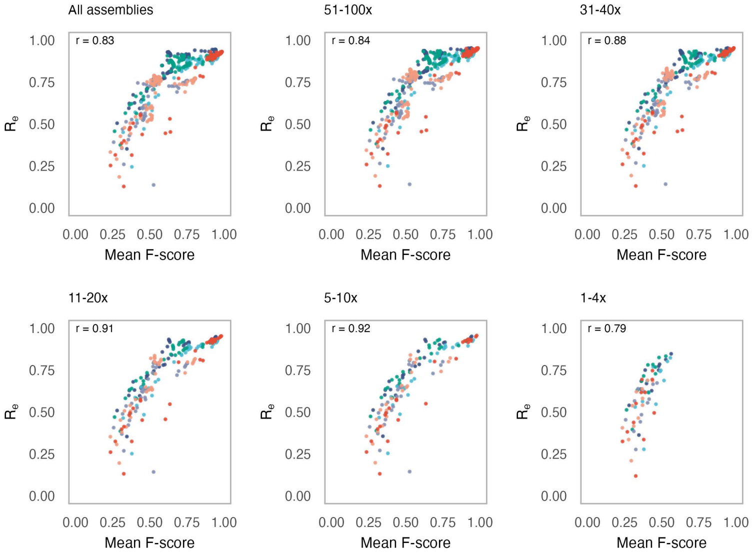

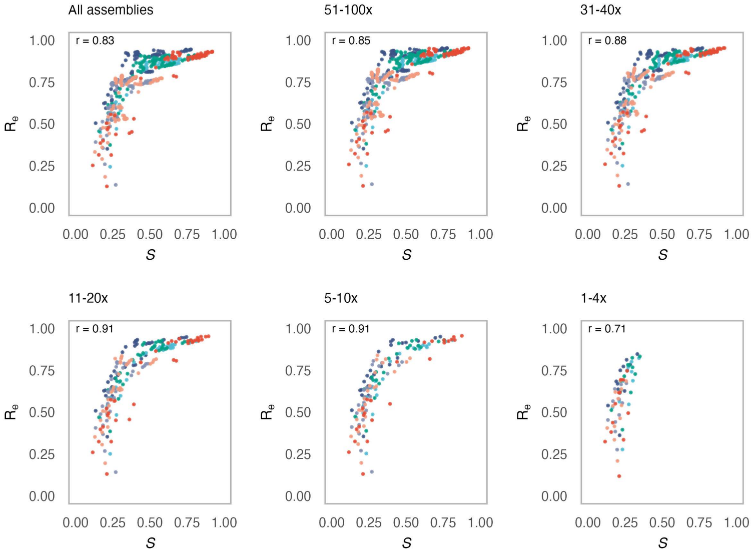

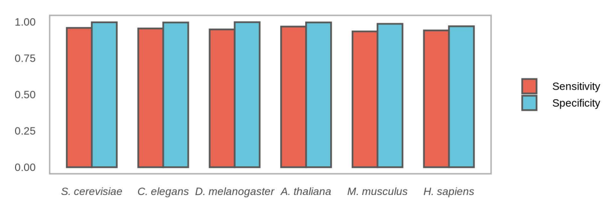
