## Supplementary Data Description for "Comprehensive Transcriptome Quality Assessment Using CATS: Reference-free and Reference-based Approaches"

**Extended Data Figures:**

**Extended data figure 1**. Distribution of CATS-rf transcript scores in assemblies generated from public RNA-seq libraries, labeled with corresponding Sequence Read Archive accession numbers. *S_t_* = CATS-rf transcript score.

**Extended data figure 2**. Distribution of CATS-rb relative exon scores across species and assemblers. *R_e_* = relative exon score. Boxplots represent the median and IQR of the distribution, with whiskers extending to ±1.5×IQR. Points denote individual assemblies.

**Extended data figure 3**. Distribution of CATS-rb relative exon scores by transcript coverage and sequencing error rate. *R_e_* = CATS-rb relative exon score. Boxplots represent the median and IQR of the distribution, with whiskers extending to ±1.5×IQR. Points denote individual assemblies.

**Extended data figure 4**. Correlation between CATS-rb relative and annotation-based exon scores in the full dataset and five subsets of simulated assemblies with decreasing coverage, as indicated in subplot titles. *R_e_* = relative exon score *, A_e_* = annotation-based exon score.

**Extended data figure 5**. Correlation between CATS-rb relative exon scores and mean transcript F-scores across the full dataset and assembly subsets with decreasing coverage, as indicated in subplot titles. *R_e_* = relative exon score.

**Extended data figure 6**. Correlation between CATS-rb relative exon scores and CATS-rf assembly scores across the full dataset and assembly subsets with decreasing coverage, as indicated in subplot titles. *R_e_* = relative exon score, *S* = CATS-rf assembly score.

**Extended data figure 7**. Performance of CATS-rb in classifying chimeric transcripts as structurally inconsistent.

**Supplementary Tables:**

**Supplementary Table 1.** Reference transcriptome assemblies used in simulating RNA-seq reads for CATS benchmarking.

**Supplementary Table 2**. Public RNA-seq libraries used in CATS benchmarking.

**Supplementary Table 3**. Reference genomes and genomic annotation used in CATS-rb benchmarking.

**Supplementary Table 4**. Species-specific CATS-rb parameters used in benchmarking.
